# Comparing machine learning models for predicting mutation status in Acute Myeloid Leukemia patients using RNA-seq data

**DOI:** 10.1101/2024.11.13.623391

**Authors:** Raíssa Silva, Cédric Riedel, Jerome Reboul, Florence Ruffle, Mélina Gallopin, Anthony Boureux, Thérèse Commes

## Abstract

Acute Myeloid Leukemia (AML) is a highly heterogeneous disease. The current AML classifications are based mainly on molecular markers, including cytogenetics features, fusion genes, and the presence or absence of mutations. In this study, we investigated mutation status in AML patients through RNA-seq data in link with differential gene expression. We applied seven machine learning algorithms to identify the presence or absence of NPM1, IDH1/IDH2, and FLT3-ITD mutations, reaching 95%, 93%, and 87% accuracy, respectively. In each case, the best performing models were complex models, suggesting highly complex biological processes at work behind AML.

## 1 Introduction

Acute Myeloid Leukemia (AML) is a hematological malignancy with high mortality. Its pathophysiology is complex and the current prognostic factors have limited predictive performance. Here, we propose a new type of analysis based on RNA-seq, k-mers, and Machine Learning (ML) algorithms to investigate biomarkers. RNA-seq data provides a great potential for identification of molecular biomakers and to target key molecular mechanisms that can drive AML pathogenesis and progression. The k-mers approach takes advantage of the fragmentation of a word (a sequence of a transcript) into multiple substrings of length k. Unlike traditional methods guided by reference annotations, k-mers approach is reference-free method. Furthermore, using k-mers can reveal alterations in certain parts of genes and also link genes together providing a new view of the investigated scenarios. K-mers offer a potential for large-scale analysis and can be particularly interesting to study mutations which frequently consist of short modifications. Coupling this data with ML algorithms allow to handle complex and non-linear links, and to integrate the complexity of the interaction between the different genes.

In this study, we compare the performance of seven classifiers: Decision Tree, K-nearest Neighbors, Logistic Regression, Neural Network, Random Forest, Support Vector Machine, and eXtreme Gradient Boosting in normalized and non-normalized data. Our objective is to apply ML algorithms to predict the prognosis of AML (for example, by predicting the presence or absence of mutations) in RNA-seq data using the k-mers approach.

## 2 Materials and Methods

This section describes the predict of the presence or absence of the NPM1, IDH1/IDH2, and FLT3-ITD mutation in AML patients.

### 2.1 Datasets

Three AML RNA-seq cohorts were used as datasets, one for training and two for testing. The training dataset is derived from the Beat-AML [1] cohort with 462 samples (access ID phs001657.v1.P1), which consists of 243 bone marrow and 219 peripheral blood samples. The test datasets are from the Leucegene [2] and Beat-AML2 [3] cohorts with 437 and 206 samples (access IDs GSE49642 and phs001657.v2.P1), respectively. The Leucegene cohort consists of 228 bone marrow and 209 peripheral blood samples. The beat-AML2 cohort consists of 118 bone marrow and 88 peripheral blood samples. Beat-AML2 samples belonging to Beat-AML patients were removed.

The cohorts used in this article were authorized for use and underwent a quality control process. We checked the quality of the raw data using fastQC version 0.11.9 [4] and MultiQC version 1.9 [5]. As a complementary quality control, we verified the sequencing protocol information and the contamination with KmerExplor [6]. After the quality control, the samples were considered good quality to be analyzed.

Henceforth, Beat-AML will be mentioned as the Training dataset, and Leucegene and Beat-AML2 as Test 1 and Test 2 datasets, respectively.

### 2.2 Features

Using k-mers has been shown to be an interesting approach to investigate several events in RNA-seq data and was used in this work as features to analyze mutation status in AML patients. To extract k-mers, we used Kmtricks [7], a tool to count k-mers efficiently in large datasets and produce a k-mer counting matrix across multiple samples. Kmtricks produced three k-mer counting matrices, one for training and two for testing (each matrix was derived from a cohort). To generate the matrices, we ran Kmtricks considering that the k-mer must have a minimum abundance of 4 (which means that the k-mer must be found at least 4 times in the sample) and be present in at least 5% of the cohort samples to be counted. The k-mer length applied was the default of the tool (size 31 nucleotides).

### 2.3 Outcome variable

AML has been linked to some mutations, including the NPM1, IDH1/IDH2 and FLT3-ITD mutations, which we chose as outcome variables. For each mutation, we classified the patients as mutated or non-mutated, making this classification binary. The number of samples per class (mutated and not mutated) and mutation type for each dataset are shown in Table Table 1.

**Table 1:** Number of mutated and non-mutated samples for NPM1, IDH1/IDH2 and FLT3-ITD mutation in Training, Test 1 and Test 2 datasets.

|  | Training |  |  | Test 1 |  |  | Test 2 |  |  |
| --- | --- | --- | --- | --- | --- | --- | --- | --- | --- |
|  | NPM1 | IDH1/IDH2 | FLT3-ITD | NPM1 | IDH1/IDH2 | FLT3-ITD | NPM1 | IDH1/IDH2 | FLT3-ITD |
| Mutated | 112 | 78 | 95 | 139 | 82 | 139 | 58 | 39 | 50 |
| Non-Mutated | 350 | 384 | 367 | 298 | 355 | 298 | 148 | 167 | 156 |

The mutation status was obtained from [Vizome](http://www.vizome.org/) website for the Training and Test 2 datasets. For the Test 1 dataset, the mutation status was obtained from [Leucegene Data website](https://data.leucegene.iric.ca/).

### 2.4 Feature selection

The k-mer counting matrix of the Training dataset consists of 366,157,215 k-mers. To reduce the data dimensionality, we developed a feature selection step. To select a k-mer, we first analyzed whether at least 70% of samples in a class are non-zero. Then, we removed the outliers (when the value was higher than the third quartile), and we calculated the mean of each class and applied the coefficient of variation to the means. If the coefficient of variation was higher or equal to 1, the k-mer was selected. The comparison of classes is given by the Equation 1 below:

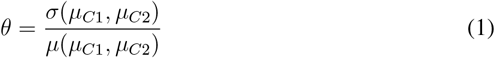

Where, *µ*_*C*1_ and *µ*_*C*2_ are the average of the samples of class 1 and class 2, respectively, *σ* is the standard deviation, and *µ* is the average. Thus, a k-mer was selected if *θ* was higher or equal to 1.

After the feature selection step, the number of features for the NPM1, IDH1/IDH2, and FLT3-ITD mutations corresponded to 26989, 774, and 717 features, respectively.

### 2.5 Machine Learning models

To predict mutation status, we selected seven Machine Learning algorithms used in other works to predict cancer [8, 9], including AML [10, 11]. We used three less complex models: Decision Tree (DT), K-nearest neighbors (KNN), and Logistic Regression (LR); and four complex models: Neural Network (NN), Random Forest (RF), Support Vector Machine (SVM), and eXtreme Gradient Boosting (XGB). We implemented the algorithms using the Scikit-Learn version 1.2.2 [12] and XGBoost version 1.7.4 [13] packages in Python, applying for each model a grid search with different parameters and a stratified cross-validation with 10-folds. For each algorithm, we built a model with original data (OD) and three normalized models with StandardScaler (SS), MinMaxScaler (MMS), and RobustScaler (RS) functions from the Scikit-Learn package.

The models were evaluated by the Receiver Operating Characteristic curve (ROC curve), Area under the ROC Curve (AUC), accuracy, sensitivity, and specificity. All the code and scripts used in this article can be found in the supplementary material.

## 3 Results and Discussion

In this section, we present the performance for Decision Tree (DT), K-Nearest Neighbors (KNN), Logistic Regression (LR), Neural Network (NN), Random Forest (RF), Support Vector Machine (SVM), and eXtreme Gradient Boosting (XGB). Each algorithm was trained with original data (OD) and normalized data with Standard Scaler (SS), Min Max Scaler (MMS), and Robust Scaler (RS).

Initially, tests with NPM1 mutation status were performed using the Beat-AML original data, 70% for training and 30% for validation, in which the Neural Network model reached 93% accuracy, 97% sensitivity, and 97% specificity. The good performance in NPM1 mutation status has led us to expand our test using the normalized data, independent datasets, and to investigate two more mutations: IDH1/IDH2 and FLT3-ITD. The initial results for the seven models are provided in the supplementary material: Table S1.

The Figure 1 shows the performance to predict NPM1 mutation status in Test 1 and Test 2 datasets. In both cases, the eXtreme Gradient Boosting trained with original data (XGB OD) achieved the best result with an AUC of 95.5% in Test 1 and 93.8% in Test 2. In addition, XGB trained with MMS normalization (XGB MMS) had a result similar to XGB OD in Test 2. Also, the XGB MMS was the second-best performance to predict in Test 1, with an AUC of 95%, and showed not to be statistically significant in difference from XGB OD when we analyzed the significance by the Wilcoxon test.

**Figure 1:**
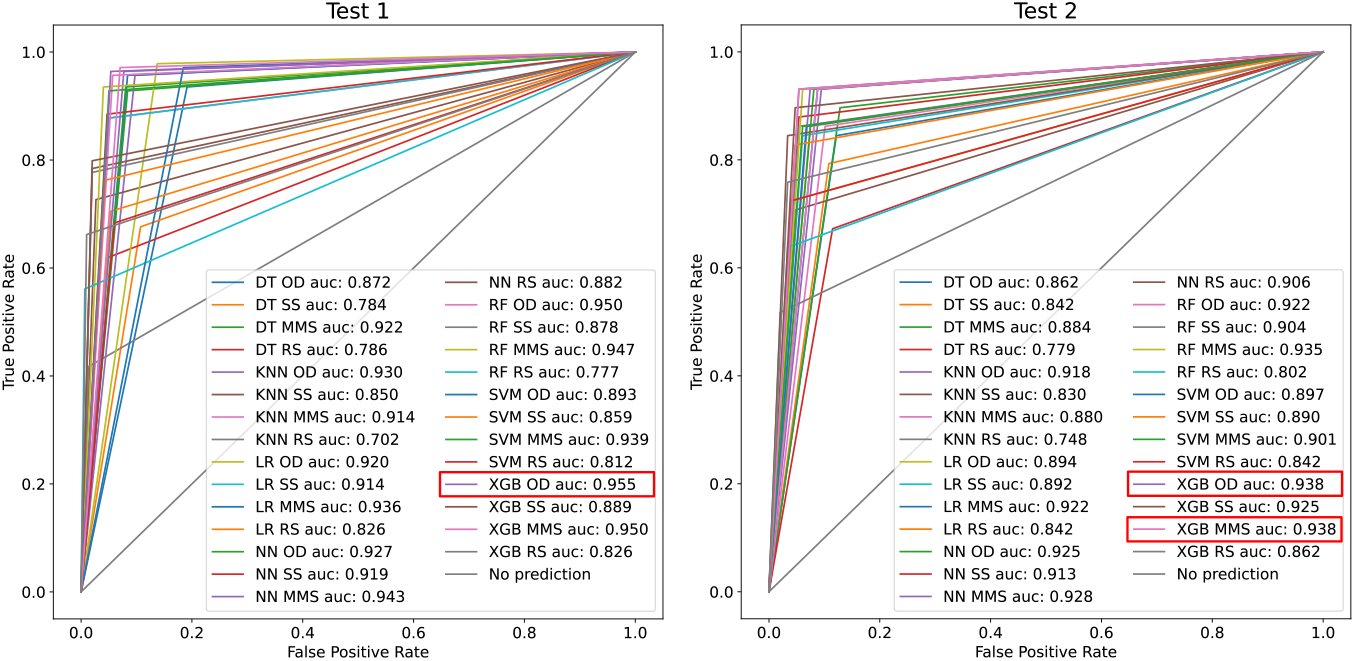
Performance for NPM1 mutation. ROC curve and AUC for models in Test 1 and Test 2 datasets. Highlight the best performers.

When analyzing the results to predict IDH1/IDH2 mutation status, Figure 2, XGB was still a good classifier. The XGB trained with SS normalization (XGB SS) achieved the best performance in both test datasets, with an AUC of 87.1% in Test 1 and 92.2% in Test 2. Although the XGB MMS and XGB OD models had similar results in the prediction of NPM1 mutation status, in the prediction of IDH1/IDH2 mutation status we did not have the same behavior between XGB SS and XGB OD. XGB OD performed less mainly in Test 1. While in Test 2, XGB OD was the second-best performance, it performed only 1% less than XGB SS, however, they showed to be statistically significant in difference by the Wilcoxon test.

**Figure 2:**
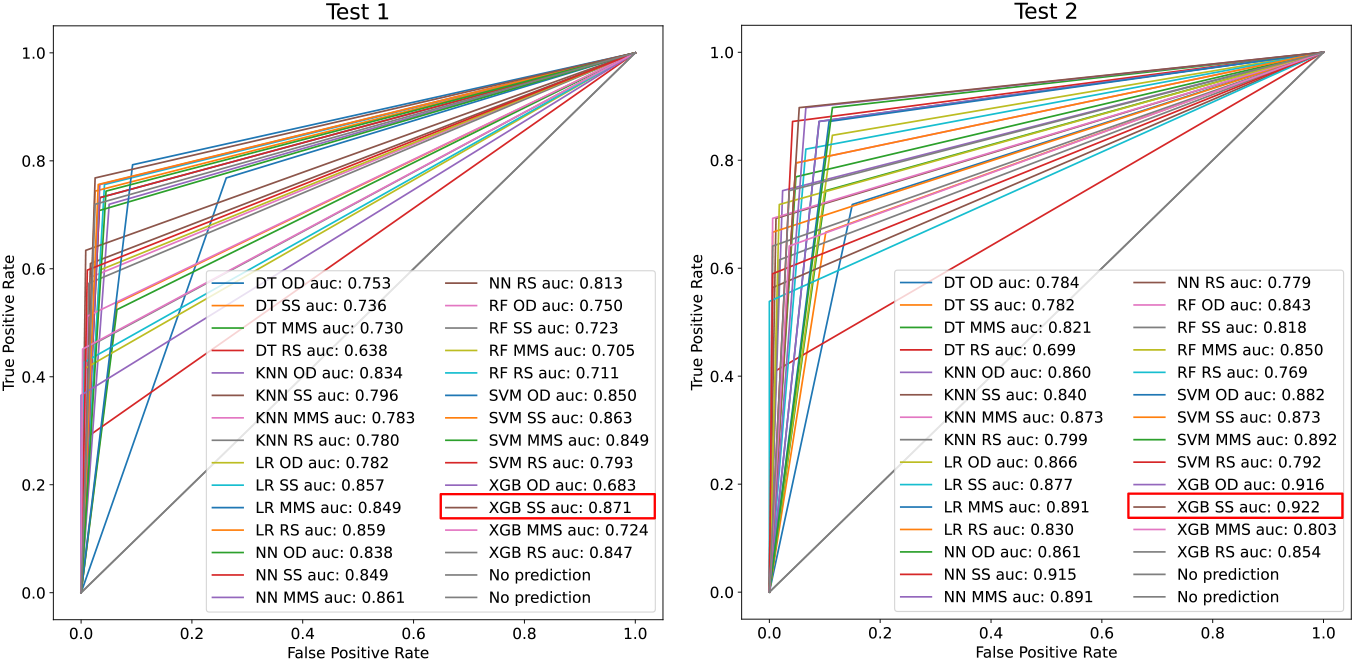
Performance for IDH1/IDH2 mutation. ROC curve and AUC for models in Test 1 and Test 2 datasets. Highlight the best performers.

The FLT3-ITD mutation status showed to be more difficult to predict. In Figure 3, the best performance was shown by RF with original data (RF OD) with an AUC of 82.7% in Test 1. The best result for Test 2 was obtained by Neural Network with MMS normalization (NN MMS), performing an AUC of 79.6%. Although the XGB did not perform well in Test 1, the XGB OD model was the second-best result in Test 2, with less than 0.01% difference. However, NN MMS and XGB OD were statistically significant in difference by the Wilcoxon test. Wilcoxon test results can be found in the supplementary material: : Table S8.

**Figure 3:**
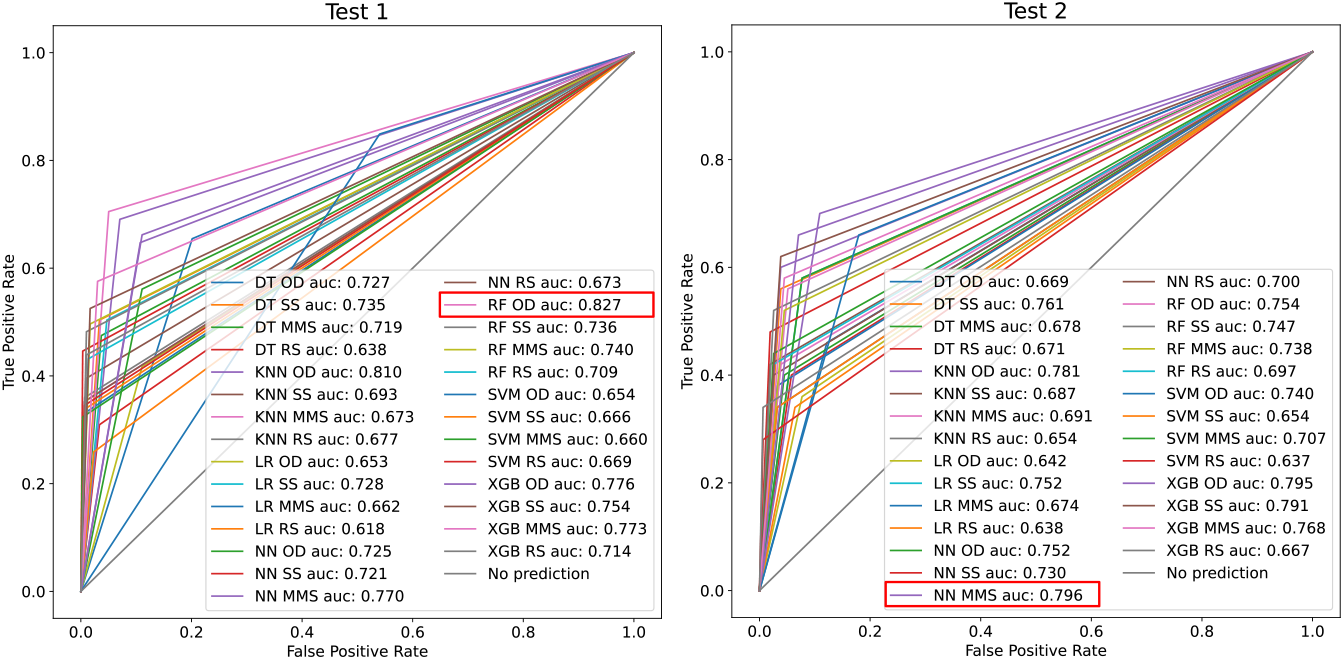
Performance for FLT3-ITD mutation. ROC curve and AUC for models in Test 1 and Test 2 datasets. Highlight the best performers.

Table 2 shows the accuracy, sensitivity, and specificity for the best models, in terms of AUC, for each set of test data by mutation. For NPM1 and IDH1/IDH2 mutations, the XGB models showed good performances, reaching more than 90% accuracy in both test datasets. While FLT3-ITD mutation reached 87% accuracy in Test 1 with RF OD and 84% of accuracy in Test 2 with NN MMS. Except for NPM1 in Test 1, the specificity was the least performed metric, on the other hand, the sensitivity was the best metric. In our case, good sensitivity means that the models can well identify patients who have the mutation. Since the main challenge is to identify a mutation, the results were promising to be further investigated.

**Table 2:** Metrics of the best machine learning algorithms when predicting NPM1, IDH1/IDH2, and FLT3-ITD mutation status.

| Mutation | Dataset | Model | Accuracy | Sensitivity | Specificity | MCC |
| --- | --- | --- | --- | --- | --- | --- |
| NPM1 | Test 1 | XGB OD | 0.951 | 0.946 | 0.964 | 0.892 |
|  | Test 2 | XGB OD | 0.941 | 0.945 | 0.931 | 0.859 |
| IDH1 / IDH2 | Test 1 | XGB SS | 0.935 | 0.974 | 0.768 | 0.781 |
|  | Test 2 | XGB SS | 0.936 | 0.946 | 0.897 | 0.806 |
| FLT3-ITD | Test 1 | RF OD | 0.871 | 0.949 | 0.705 | 0.696 |
|  | Test 2 | NN MMS | 0.844 | 0.891 | 0.700 | 0.583 |

Because of the imbalanced nature of the data, we measured the performance by Matthew’s correlation coefficient (MCC). Furthermore, we also used Synthetic Minority Oversampling Technique (SMOTE) to limit classes disproportion, however the results was similar when comparing to the original data (see supplementary material: Additional information 1).

Metrics for all models can be found in the supplementary material: Tables S2-S7.

## 4 Final considerations

We applied seven machine learning algorithms to predict mutation status in AML with normalized and non-standard data. XGB demonstrated good performance to predict NPM1 and IDH1/IDH2 mutation status, while the FLT3-ITD mutation required different models for each test dataset. In an additional analysis of our selected features, NPM1, IDH1/IDH2, and FLT3-ITD mutations were correlated to 164, 44, and 28 annotated genes, respectively. Mutations in NPM1 are the most common alterations in AML with a recurrently mutated position, frequently associated with a favorable prognosis [14]. From the biological point of view, the mutation has a clear impact on leukemia cells and it is not surprising to obtain the best prediction for this group of patients as well as the higher number of differential selected genes. These results will guide us to explore further the predictions in the diagnosis and prognosis of AML, as well as to analyze the genes and pathways involved. Furthermore, these results encourages us to study whether the impact of the mutation on other genes is proportional to the performance of the models. In this end, we plan to use eXplainable Artificial Intelligence methods to analyze the correlation of k-mers and explore the link between the genes involved.

## Availability of data and software code

Our codes, scripts, supplementary material, and more detail about the machine learning algorithm configuration parameters values used are available at https://osf.io/cw9td/.

