## Supplementary material for "Comparing machine learning models for predicting mutation status in Acute Myeloid Leukemia patients using RNA-seq data"

**Supplementary Table 1.** Accuracy, Kappa Sensibility, and Specificity for dataset validation with Beat-AML using original data (70% to train, 30% to test).

| Model | Accuracy | Sensitivity | Specificity |
|---|---|---|---|
| Decision Tree (DT) | 0.827 | 0.878 | 0.656 |
| K-nearest neighbors (KNN) | 0.870 | 0.933 | 0.676 |
| Logistic Regression (LR) | 0.913 | 0.932 | 0.857 |
| Neural Network (NN) | **0.935** | 0.971 | **0.971** |
| Random Forest (RF) | 0.913 | **0.973** | 0.64 |
| Support Vector Machine (SVM) | 0.884 | 0.960 | 0.684 |
| eXtreme Gradient Boosting (XGB) | 0.928 | 0.933 | 0.909 |

**Supplementary Table 2.** Performance to predict NPM1 mutation status in Test 1.

| Model | AUC | Accuracy | Sensitivity | Specificity | MCC |
|---|---|---|---|---|---|
| Decision tree OD | 0.871988 | 0.84897 | 0.808725 | 0.935252 | 0.7003 |
| Decision tree SS | 0.784438 | 0.823799 | 0.892617 | 0.676259 | 0.584882 |
| Decision tree MMS | 0.922082 | 0.919908 | 0.916107 | 0.928058 | 0.82298 |
| Decision tree RS | 0.785863 | 0.846682 | 0.95302 | 0.618705 | 0.633852 |
| K-nearest neighbors OD | 0.92976 | 0.919908 | 0.902685 | 0.956835 | 0.828795 |
| K-nearest neighbors SS | 0.849887 | 0.894737 | 0.973154 | 0.726619 | 0.753221 |
| K-nearest neighbors MMS | 0.913681 | 0.926773 | 0.949664 | 0.877698 | 0.830597 |
| K-nearest neighbors RS | 0.701922 | 0.805492 | 0.986577 | 0.417266 | 0.539035 |
| Logistic Regression OD | 0.920417 | 0.899314 | 0.862416 | **0.978417** | 0.797723 |
| Logistic Regression SS | 0.913681 | 0.926773 | 0.949664 | 0.877698 | 0.830597 |
| Logistic Regression MMS | 0.936471 | 0.929062 | 0.916107 | 0.956835 | 0.846195 |
| Logistic Regression RS | 0.825672 | 0.869565 | 0.946309 | 0.705036 | 0.690832 |
| Neural Network OD | 0.927357 | 0.924485 | 0.919463 | 0.935252 | 0.833265 |
| Neural Network SS | 0.918956 | 0.93135 | 0.95302 | 0.884892 | 0.841188 |
| Neural Network MMS | 0.943424 | 0.935927 | 0.922819 | 0.964029 | 0.860866 |
| Neural Network RS | 0.882019 | 0.91762 | 0.979866 | 0.784173 | 0.808079 |
| Random Forest OD | 0.949894 | 0.947368 | 0.942953 | 0.956835 | 0.88261 |
| Random Forest SS | 0.878422 | 0.915332 | 0.979866 | 0.776978 | 0.802728 |
| Random Forest MMS | 0.947492 | 0.951945 | 0.959732 | 0.935252 | 0.88997 |
| Random Forest RS | 0.77722 | 0.855835 | **0.993289** | 0.561151 | 0.667714 |
| Support Vector Machine OD | 0.89333 | 0.864989 | 0.815436 | 0.971223 | 0.739055 |
| Support Vector Machine SS | 0.859483 | 0.894737 | 0.956376 | 0.76259 | 0.752206 |
| Support Vector Machine MMS | 0.938861 | 0.942792 | 0.949664 | 0.928058 | 0.869676 |
| Support Vector Machine RS | 0.811525 | 0.858124 | 0.939597 | 0.683453 | 0.662715 |
| Extreme Gradient Boosting OD | **0.955169** | **0.951945** | 0.946309 | 0.964029 | **0.892959** |
| Extreme Gradient Boosting SS | 0.889213 | 0.922197 | 0.979866 | 0.798561 | 0.818777 |
| Extreme Gradient Boosting MMS | 0.950377 | 0.942792 | 0.92953 | 0.971223 | 0.8756 |
| Extreme Gradient Boosting RS | 0.825902 | 0.885584 | 0.989933 | 0.661871 | 0.735965 |

**Supplementary Table 3.** Performance to predict NPM1 mutation status in Test 2.

| Model | AUC | Accuracy | Sensitivity | Specificity | MCC |
|---|---|---|---|---|---|
| Decision tree OD | 0.861603 | 0.868932 | 0.878378 | 0.844828 | 0.694324 |
| Decision tree SS | 0.842498 | 0.864078 | 0.891892 | 0.793103 | 0.671669 |
| Decision tree MMS | 0.884087 | 0.878641 | 0.871622 | 0.896552 | 0.726956 |
| Decision tree RS | 0.778774 | 0.825243 | 0.885135 | 0.672414 | 0.563622 |
| K-nearest neighbors OD | 0.91822 | 0.912621 | 0.905405 | **0.931034** | 0.799992 |
| K-nearest neighbors SS | 0.8298 | 0.883495 | 0.952703 | 0.706897 | 0.701739 |
| K-nearest neighbors MMS | 0.880359 | 0.88835 | 0.898649 | 0.862069 | 0.736211 |
| K-nearest neighbors RS | 0.748486 | 0.849515 | **0.97973** | 0.517241 | 0.609391 |
| Logistic Regression OD | 0.893872 | 0.907767 | 0.925676 | 0.862069 | 0.776035 |
| Logistic Regression SS | 0.892008 | 0.912621 | 0.939189 | 0.844828 | 0.784017 |
| Logistic Regression MMS | 0.921598 | 0.917476 | 0.912162 | **0.931034** | 0.809522 |
| Logistic Regression RS | 0.841799 | 0.893204 | 0.959459 | 0.724138 | 0.727271 |
| Neural Network OD | 0.924977 | 0.92233 | 0.918919 | **0.931034** | 0.819226 |
| Neural Network SS | 0.912628 | 0.927184 | 0.945946 | 0.87931 | 0.821011 |
| Neural Network MMS | 0.928355 | 0.927184 | 0.925676 | **0.931034** | 0.829111 |
| Neural Network RS | 0.905522 | 0.932039 | 0.966216 | 0.844828 | 0.829412 |
| Random Forest OD | 0.921598 | 0.917476 | 0.912162 | **0.931034** | 0.809522 |
| Random Forest SS | 0.904007 | 0.92233 | 0.945946 | 0.862069 | 0.808015 |
| Random Forest MMS | 0.935112 | 0.936893 | 0.939189 | **0.931034** | 0.849449 |
| Random Forest RS | 0.802074 | 0.873786 | 0.966216 | 0.637931 | 0.674437 |
| Support Vector Machine OD | 0.897251 | 0.912621 | 0.932432 | 0.862069 | 0.78648 |
| Support Vector Machine SS | 0.890144 | 0.917476 | 0.952703 | 0.827586 | 0.793287 |
| Support Vector Machine MMS | 0.900629 | 0.917476 | 0.939189 | 0.862069 | 0.797136 |
| Support Vector Machine RS | 0.841799 | 0.893204 | 0.959459 | 0.724138 | 0.727271 |
| Extreme Gradient Boosting OD | **0.93849** | **0.941748** | 0.945946 | **0.931034** | **0.859919** |
| Extreme Gradient Boosting SS | 0.924627 | 0.936893 | 0.952703 | 0.896552 | 0.844886 |
| Extreme Gradient Boosting MMS | **0.93849** | **0.941748** | 0.945946 | **0.931034** | **0.859919** |
| Extreme Gradient Boosting RS | 0.862418 | 0.907767 | 0.966216 | 0.758621 | 0.765662 |

**Supplementary Table 4.** Performance to predict IDH1/IDH2 mutation status in Test 1.

| Model | AUC | Accuracy | Sensitivity | Specificity | MCC |
|---|---|---|---|---|---|
| Decision tree OD | 0.75316 | 0.743707 | 0.738028 | 0.768293 | 0.412602 |
| Decision tree SS | 0.735898 | 0.860412 | 0.935211 | 0.536585 | 0.511255 |
| Decision tree MMS | 0.729801 | 0.858124 | 0.935211 | 0.52439 | 0.501122 |
| Decision tree RS | 0.637891 | 0.853547 | 0.983099 | 0.292683 | 0.425822 |
| K-nearest neighbors OD | 0.834404 | 0.906178 | 0.949296 | 0.719512 | 0.685371 |
| K-nearest neighbors SS | 0.796427 | 0.913043 | 0.983099 | 0.609756 | 0.692489 |
| K-nearest neighbors MMS | 0.782824 | 0.906178 | 0.980282 | 0.585366 | 0.665816 |
| K-nearest neighbors RS | 0.779543 | 0.908467 | 0.985915 | 0.573171 | 0.674168 |
| Logistic Regression OD | 0.781879 | 0.897025 | 0.966197 | 0.597561 | 0.635119 |
| Logistic Regression SS | 0.856922 | 0.919908 | 0.957746 | 0.756098 | 0.731523 |
| Logistic Regression MMS | 0.848952 | 0.922197 | 0.966197 | 0.731707 | 0.734521 |
| Logistic Regression RS | 0.859275 | 0.93135 | 0.974648 | 0.743902 | 0.764885 |
| Neural Network OD | 0.838166 | 0.919908 | 0.969014 | 0.707317 | 0.724156 |
| Neural Network SS | 0.848952 | 0.922197 | 0.966197 | 0.731707 | 0.734521 |
| Neural Network MMS | 0.861147 | 0.926773 | 0.966197 | 0.756098 | 0.751911 |
| Neural Network RS | 0.812848 | 0.924485 | 0.991549 | 0.634146 | 0.736497 |
| Random Forest OD | 0.750464 | 0.899314 | 0.988732 | 0.512195 | 0.637278 |
| Random Forest SS | 0.722793 | 0.892449 | 0.994366 | 0.45122 | 0.610209 |
| Random Forest MMS | 0.7045 | 0.885584 | 0.994366 | 0.414634 | 0.580793 |
| Random Forest RS | 0.710598 | 0.887872 | 0.994366 | 0.426829 | 0.590709 |
| Support Vector Machine OD | 0.849863 | 0.885584 | 0.907042 | **0.792683** | 0.654992 |
| Support Vector Machine SS | 0.862556 | 0.929062 | 0.969014 | 0.756098 | 0.758951 |
| Support Vector Machine MMS | 0.849416 | 0.915332 | 0.95493 | 0.743902 | 0.716139 |
| Support Vector Machine RS | 0.793147 | 0.915332 | 0.988732 | 0.597561 | 0.701185 |
| Extreme Gradient Boosting OD | 0.682927 | 0.881007 | **1** | 0.365854 | 0.564899 |
| Extreme Gradient Boosting SS | **0.87147** | **0.935927** | 0.974648 | 0.768293 | **0.78192** |
| Extreme Gradient Boosting MMS | 0.724201 | 0.894737 | 0.997183 | 0.45122 | 0.621314 |
| Extreme Gradient Boosting RS | 0.84708 | 0.926773 | 0.974648 | 0.719512 | 0.747675 |

**Supplementary Table 5.** Performance to predict IDH1/IDH2 mutation status in Test 2.

| Model | AUC | Accuracy | Sensitivity | Specificity | MCC |
|---|---|---|---|---|---|
| Decision tree OD | 0.784124 | 0.825243 | 0.850299 | 0.717949 | 0.509266 |
| Decision tree SS | 0.782435 | 0.854369 | 0.898204 | 0.666667 | 0.544517 |
| Decision tree MMS | 0.820897 | 0.868932 | 0.898204 | 0.74359 | 0.603736 |
| Decision tree RS | 0.69914 | 0.878641 | 0.988024 | 0.410256 | 0.552541 |
| K-nearest neighbors OD | 0.859819 | 0.932039 | 0.976048 | 0.74359 | 0.768642 |
| K-nearest neighbors SS | 0.840166 | 0.932039 | 0.988024 | 0.692308 | 0.766347 |
| K-nearest neighbors MMS | 0.873484 | 0.92233 | 0.952096 | 0.794872 | 0.746968 |
| K-nearest neighbors RS | 0.79871 | 0.912621 | 0.982036 | 0.615385 | 0.693525 |
| Logistic Regression OD | 0.866191 | 0.878641 | 0.886228 | 0.846154 | 0.660489 |
| Logistic Regression SS | 0.877322 | 0.912621 | 0.934132 | 0.820513 | 0.727453 |
| Logistic Regression MMS | 0.890987 | 0.902913 | 0.91018 | 0.871795 | 0.719507 |
| Logistic Regression RS | 0.830339 | 0.932039 | 0.994012 | 0.666667 | 0.766959 |
| Neural Network OD | 0.860663 | 0.917476 | 0.952096 | 0.769231 | 0.728578 |
| Neural Network SS | 0.914939 | **0.941748** | 0.958084 | 0.871795 | **0.814275** |
| Neural Network MMS | 0.890987 | 0.902913 | 0.91018 | 0.871795 | 0.719507 |
| Neural Network RS | 0.779057 | 0.912621 | 0.994012 | 0.564103 | 0.694264 |
| Random Forest OD | 0.84316 | 0.936893 | 0.994012 | 0.692308 | 0.784563 |
| Random Forest SS | 0.817519 | 0.927184 | 0.994012 | 0.641026 | 0.749147 |
| Random Forest MMS | 0.849992 | 0.932039 | 0.982036 | 0.717949 | 0.766971 |
| Random Forest RS | 0.769231 | 0.912621 | **1** | 0.538462 | 0.697188 |
| Support Vector Machine OD | 0.882005 | 0.88835 | 0.892216 | 0.871795 | 0.689014 |
| Support Vector Machine SS | 0.873484 | 0.92233 | 0.952096 | 0.794872 | 0.746968 |
| Support Vector Machine MMS | 0.891832 | 0.88835 | 0.886228 | **0.897436** | 0.698074 |
| Support Vector Machine RS | 0.791878 | 0.917476 | 0.994012 | 0.589744 | 0.712821 |
| Extreme Gradient Boosting OD | 0.915784 | 0.927184 | 0.934132 | **0.897436** | 0.782257 |
| Extreme Gradient Boosting SS | **0.921772** | 0.936893 | 0.946108 | **0.897436** | 0.806333 |
| Extreme Gradient Boosting MMS | 0.802549 | 0.902913 | 0.964072 | 0.641026 | 0.663004 |
| Extreme Gradient Boosting RS | 0.853831 | 0.92233 | 0.964072 | 0.74359 | 0.738217 |

**Supplementary Table 6.** Performance to predict IFT3-ITD mutation status in Test 1.

| Model | AUC | Accuracy | Sensitivity | Specificity | MCC |
|---|---|---|---|---|---|
| Decision tree OD | 0.726667 | 0.75286 | 0.798658 | 0.654676 | 0.443979 |
| Decision tree SS | 0.73502 | 0.819222 | 0.966443 | 0.503597 | 0.566072 |
| Decision tree MMS | 0.718954 | 0.80778 | 0.963087 | 0.47482 | 0.535305 |
| Decision tree RS | 0.637898 | 0.757437 | 0.966443 | 0.309353 | 0.393459 |
| K-nearest neighbors OD | 0.810089 | 0.853547 | 0.92953 | 0.690647 | 0.652325 |
| K-nearest neighbors SS | 0.692808 | 0.800915 | 0.989933 | 0.395683 | 0.529343 |
| K-nearest neighbors MMS | 0.672903 | 0.789474 | 0.993289 | 0.352518 | 0.501614 |
| K-nearest neighbors RS | 0.6765 | 0.791762 | 0.993289 | 0.359712 | 0.507761 |
| Logistic Regression OD | 0.653264 | 0.73913 | 0.889262 | 0.417266 | 0.351582 |
| Logistic Regression SS | 0.728309 | 0.810069 | 0.95302 | 0.503597 | 0.539686 |
| Logistic Regression MMS | 0.662112 | 0.782609 | 0.993289 | 0.330935 | 0.482909 |
| Logistic Regression RS | 0.617751 | 0.748284 | 0.97651 | 0.258993 | 0.368239 |
| Neural Network OD | 0.725206 | 0.784897 | 0.889262 | 0.561151 | 0.481899 |
| Neural Network SS | 0.721344 | 0.82151 | **0.996644** | 0.446043 | 0.586958 |
| Neural Network MMS | 0.77005 | 0.814645 | 0.892617 | 0.647482 | 0.56073 |
| Neural Network RS | 0.672903 | 0.789474 | 0.993289 | 0.352518 | 0.501614 |
| Random Forest OD | **0.82735** | **0.871854** | 0.949664 | 0.705036 | **0.69638** |
| Random Forest SS | 0.735974 | 0.828375 | 0.989933 | 0.482014 | 0.599276 |
| Random Forest MMS | 0.739812 | 0.828375 | 0.983221 | 0.496403 | 0.59559 |
| Random Forest RS | 0.709116 | 0.810069 | 0.986577 | 0.431655 | 0.55092 |
| Support Vector Machine OD | 0.654326 | 0.583524 | 0.459732 | **0.848921** | 0.299195 |
| Support Vector Machine SS | 0.665709 | 0.784897 | 0.993289 | 0.338129 | 0.48919 |
| Support Vector Machine MMS | 0.660193 | 0.782609 | **0.996644** | 0.323741 | 0.486207 |
| Support Vector Machine RS | 0.669306 | 0.787185 | 0.993289 | 0.345324 | 0.495425 |
| Extreme Gradient Boosting OD | 0.775566 | 0.816934 | 0.889262 | 0.661871 | 0.567988 |
| Extreme Gradient Boosting SS | 0.754201 | 0.837529 | 0.983221 | 0.52518 | 0.61834 |
| Extreme Gradient Boosting MMS | 0.772669 | 0.844394 | 0.969799 | 0.57554 | 0.630662 |
| Extreme Gradient Boosting RS | 0.714391 | 0.814645 | 0.989933 | 0.438849 | 0.564817 |

**Supplementary Table 7.** Performance to predict IFT3-ITD mutation status in Test 2.

| Model | AUC | Accuracy | Sensitivity | Specificity | MCC |
|---|---|---|---|---|---|
| Decision tree OD | 0.669103 | 0.776699 | 0.878205 | 0.46 | 0.359899 |
| Decision tree SS | 0.760769 | 0.864078 | 0.961538 | 0.56 | 0.602324 |
| Decision tree MMS | 0.677564 | 0.820388 | 0.955128 | 0.4 | 0.451153 |
| Decision tree RS | 0.671154 | 0.81068 | 0.942308 | 0.4 | 0.421967 |
| K-nearest neighbors OD | 0.780769 | 0.873786 | 0.961538 | 0.6 | 0.633944 |
| K-nearest neighbors SS | 0.687179 | 0.834951 | 0.974359 | 0.4 | 0.500258 |
| K-nearest neighbors MMS | 0.690769 | 0.830097 | 0.961538 | 0.42 | 0.484705 |
| K-nearest neighbors RS | 0.653974 | 0.815534 | 0.967949 | 0.34 | 0.42747 |
| Logistic Regression OD | 0.641538 | 0.786408 | 0.923077 | 0.36 | 0.34406 |
| Logistic Regression SS | 0.751538 | 0.839806 | 0.923077 | 0.58 | 0.540192 |
| Logistic Regression MMS | 0.673974 | 0.825243 | 0.967949 | 0.38 | 0.464966 |
| Logistic Regression RS | 0.637949 | 0.791262 | 0.935897 | 0.34 | 0.350499 |
| Neural Network OD | 0.751538 | 0.839806 | 0.923077 | 0.58 | 0.540192 |
| Neural Network SS | 0.730385 | 0.859223 | 0.980769 | 0.48 | 0.585359 |
| Neural Network MMS | **0.795513** | 0.84466 | 0.891026 | **0.7** | 0.583299 |
| Neural Network RS | 0.700385 | 0.84466 | 0.980769 | 0.42 | 0.53555 |
| Random Forest OD | 0.754359 | 0.854369 | 0.948718 | 0.56 | 0.574313 |
| Random Forest SS | 0.747179 | 0.864078 | 0.974359 | 0.52 | 0.600859 |
| Random Forest MMS | 0.737564 | 0.849515 | 0.955128 | 0.52 | 0.555364 |
| Random Forest RS | 0.697179 | 0.839806 | 0.974359 | 0.42 | 0.517761 |
| Support Vector Machine OD | 0.740256 | 0.781553 | 0.820513 | 0.66 | 0.451236 |
| Support Vector Machine SS | 0.653974 | 0.815534 | 0.967949 | 0.34 | 0.42747 |
| Support Vector Machine MMS | 0.707179 | 0.84466 | 0.974359 | 0.44 | 0.534935 |
| Support Vector Machine RS | 0.636795 | 0.820388 | 0.99359 | 0.28 | 0.451424 |
| Extreme Gradient Boosting OD | 0.794744 | 0.864078 | 0.929487 | 0.66 | 0.616649 |
| Extreme Gradient Boosting SS | 0.790769 | **0.878641** | 0.961538 | 0.62 | **0.649504** |
| Extreme Gradient Boosting MMS | 0.767564 | 0.864078 | 0.955128 | 0.58 | 0.604129 |
| Extreme Gradient Boosting RS | 0.666795 | 0.834951 | **0.99359** | 0.34 | 0.50646 |

**Supplementary Table 8.** Wilcoxon test.

| Mutation | Dataset | Models | Statistic | P-value |
|---|---|---|---|---|
| NPM1 | Test 1 | XGB OD and XGB MMS | -0.4250 | 0.6709 |
| | Test 2 | XGB OD and XGB MMS | 1.5 | 1.0 |
| IDH1 / IDH2 | Test 2 | XGB OD and XGB SS | 0.0 | 1.182e-11 |
| FLT3-ITD | Test 2 | NN_MMS and XGB OD | 0.0 | 1.085e-20 |

**Additional information 1.** Because mutation prediction is an unbalanced classification, we did tests using Synthetic Minority Oversampling Technique (SMOTE) by imbalanced-learn Python library to balance the classes. The results using the SMOTE technique did not have a high impact on performance. Thus, we chose to present in the paper the results with unbalanced data because it represents the reality of the biological data. Below are presented the results for the best model with imbalanced and balanced data:

| Imbalanced data | | | | | | |
|---|---|---|---|---|---|---|
| Mutation | Dataset | Model | AUC | Accuracy | Sens | Spec |
| NPM1 | Test 1 | XGB OD | **0.955** | **0.951** | **0.946** | 0.964 |
| | Test 2 | XGB OD | **0.938** | **0.941** | **0.945** | **0.931** |
| IDH1 / IDH2 | Test 1 | XGB SS | **0.871** | **0.935** | **0.974** | **0.768** |
| | Test 2 | XGB SS | 0.921 | 0.936 | 0.946 | **0.897** |
| FLT3-ITD | Test 1 | RF OD | **0.827** | **0.871** | **0.949** | 0.705 |
| | Test 2 | NN MMS | **0.795** | 0.844 | 0.891 | **0.7** |
| Balanced data (SMOTE) | | | | | | |
| Mutation | Dataset | Model | AUC | Accuracy | Sens | Spec |
| NPM1 | Test 1 | RF OD | 0.950 | 0.942 | 0.929 | **0.971** |
| | Test 2 | XGB OD | **0.938** | **0.941** | **0.945** | **0.931** |
| IDH1 / IDH2 | Test 1 | RF MMS | 0.864 | 0.924 | 0.960 | **0.768** |
| | Test 2 | XGB OD | **0.925** | **0.941** | **0.952** | **0.897** |
| FLT3-ITD | Test 1 | RF OD | 0.826 | 0.867 | 0.939 | **0.712** |
| | Test 2 | XGB OD | 0.794 | **0.864** | **0.929** | 0.66 |

**Additional information 2.** For each mutation, we applied STAR 2.7.8a [1] to map the selected features (k-mers) to a reference human genome, GRCh38 assembly. Afterwards, we used Samtools 1.11 [2] to generate flexible alignment formats, SAM, BED, and BAM files. Next, we implemented a script in R using the Ensembl REST API [3] to request the genes annotation for each k-mer using the SAM and BAM files.

Annotated genes related to 26,989 selected features (k-mers) for NPM1 mutation:
NBL1, DLGAP3, CCDC24, SLC6A9, DYNLT5, KRTCAP2, TRIM46, LY9, POU2F1, CRYZL2P-SEC16B, SEC16B, CRYZL2P, IPO9-AS1, NAV1, PFKFB2, CD34, TRIB2, RASGRP3, MEIS1, MEIS1-AS2, M1AP, IGKV2-23, GYPC, FTCDNL1, MREG, USP40, RAMP1, VGLL4, MYRIP, EIF1B-AS1, CRYBG3, CD200, TRH, IGSF10, MED12L, IL12A-AS1, C3orf80, IFT80, SMC4, LMLN, ZNF595, IDUA, SPON2, PGRMC2, OTULINL, ANKH, RETREG1, RETREG1-AS1, ANXA2R-OT1, MARVELD2, OCLN, LINC01455, TSLP, LINC02147, LINC02208, FOXC1, H2BC15, PREP, RPS6KA2, TSPAN13, SNX10, SNX10-AS1, LINC02981, HOXA2, HOXA3, HOXA-AS2, HOXA4, HOXA-AS3, HOXA5, HOXA6, HOXA7, HOXA9, HOXA10-AS, HOXA10, ABCB1, GNG11, CCDC136, ADAM7-AS1, ADAMDEC1, BAALC, BAALC-AS1, TMEM65, ADGRB1, MLLT3, GGTA1, PBX3, SARDH, ST8SIA6, ST8SIA6-AS1, SKIDA1, ARHGAP22, ANK3, LINC01475, NKX2-3, GSTO1, CAT, PIWIL4, PIWIL4-AS1, PDGFD, SIAE, ESAM, CACNA2D4, CPNE8, PRICKLE1, PPHLN1, POU6F1, SMAGP, DAZAP2, SDS, SDSL, UGGT2, LINC01500, BRF1, ATP10A, GOLGA8M, OIP5-AS1, MYEF2, DMXL2, FBXO22, NRG4, HOMER2, LINC01569, IQCK, GPRC5B, ZNF688, IRX3, CRNDE, IRX5, WWOX, SERPINF1, FBXL20, ACLY, HOXB2, HOXB-AS1, HOXB3, HOXB-AS3, HOXB-AS2, HOXB4, HOXB5, HOXB6, HOXB7, TOM1L1, RENO1, LINC03048, BAHCC1, MTCL1, LDLRAD4, SETBP1, TMIGD2, ADGRE1, ADAMTS10, YJU2B, ZNF626, SCN1B, LINC02910, APP, LINC01637, LINC00899, PRR34, PRR34-AS1, MIRLET7BHG, PHKA2-AS1, COL4A5, FHL1

Annotated genes related to 774 selected features (k-mers) for IDH1/IDH2 mutation:
HENMT1, C1orf21, LAX1, PTH2R, SP140, HRH1, NUDT16-DT, NUDT16L2P, ANAPC13, ZNF518B, CCSER1, RAP1GDS1, SLC9A3, SLC9A3-AS1, C2, TNXB, GPSM3, DST, SRSF12, DDO, ASIC3, FAM86B3P, CA3-AS1, CA2, ANKRD18B, DNAJC1, PAOX, MIR210HG, MRPL23, TPCN1, CAPN3, CLCN7, ROGDI, SERPINF1, GOSR2, ERFL, ARHGEF1, CKM, EPS8L1, CD93, PPM1F, PPM1F-AS1, LRRC75B, F8

Annotated genes related to 717 selected features (k-mers) for IFT3-ITD mutation:
SHISAL2A, CFH, LPIN1, MAP4K3-DT, TRAK2, STRADB, MAPKAPK3, SCHIP1, IQCJ-SCHIP1, FAM47E, FAM47E-STBD1, HOXA-AS3, HOXA6, HOXA3, BMP1, PTBP3, SOCS2-AS1, SOCS2, VSIG10, STON2, GOLGA8B, LYRM1, ADGRG1, TRIM16, MSI2, TBC1D16, LINC02940, APOL4
